# Baseline State for Pulmonary Vasculature with Pulmonary Arterial Hypertension: Effect of Geometric Remodeling and Metabolic Shift

**DOI:** 10.1101/2025.04.30.651520

**Authors:** Haritha N. Mullagura, Hamidreza Gharahi, C. Alberto Figueroa, Seungik Baek

**Affiliations:** Department of Mechanical Engineering, Michigan State University, East Lansing, MI; Section of Vascular Surgery, Department of Surgery, University of Michigan, Ann Arbor, MI; Department of Biomedical Engineering, University of Michigan, Ann Arbor, MI

**Keywords:** Hemodynamics, Mechanical homeostasis, Metabolic shift, Pulmonary hypertension

## Abstract

Pulmonary arterial hypertension (PAH) is a complex disease characterized by chronically elevated pulmonary arterial pressure, with early onset and progression linked to structural, metabolic and morphological changes in the pulmonary vasculature. Understanding the interplay between hemodynamics and arterial wall mechanics is essential to capture the pathology of the distal vasculature in PAH. This study aims to develop a data-driven framework that establishes a baseline state of PAH vasculature, incorporating key features of arterial wall constituents, geometry, and their interaction with PAH-specific hemodynamics. The model also explores changes in the metabolic energy costs of the arterial vasculature and hypothesize the most plausible metabolic costs based on PAH arterial wall energy consumption. Illustrative examples of symmetrically bifurcating arterial trees are used to establish baseline characteristics of PAH-affected pulmonary arteries. We establish the baseline state for PAH vasculature and compared it with healthy homeostatic vasculature in terms of arterial mechanics, morphometry, and pulsatile hemodynamics. This framework provides a representative computational model for advanced studies on PAH treatments and lays the groundwork for future pathophysiological modeling of the disease.

## 1. Introduction

Pulmonary arterial hypertension (PAH) is a multifaceted vascular disease marked by progressive vascular remodeling, resulting in narrowing of the pulmonary arteries and increased pulmonary vascular resistance (PVR). These changes cause in elevated afterload on the right heart, leading to right heart failure if left untreated (Thenappan et al. 2018b). These pathological changes include endothelial cell (EC) proliferation, smooth muscle cell (SMCs) hypertrophy, and excessive collagen deposition in the extracellular matrix (ECM) (Chazova et al. 1995; Thenappan et al. 2018a). These factors contribute to thickened and stiffened arterial walls, fundamentally altering the mechanical behavior of the pulmonary vasculature and pulmonary hemodynamic circulation. Clinical imaging is, however, limited in their ability to detect the vascular features of small vessels and human tissue samples from PAH patients are scarce (Shahin et al. 2022). To bridge this gap, computational modeling of pulmonary arterial vasculature has emerged as a valuable tool, offering insight into the relationship between pulmonary geometrical features and hemodynamic function, and complex vascular remodeling processes, despite the limited available experimental data (Hunter et al. 2010b; Qureshi and Hill 2015; Ebrahimi et al. 2022; Szafron et al. 2023).

More recent efforts have focused on PAH-specific models that incorporate stress-mediated growth and remodeling (G&R), such as those that simulate changes in pulmonary artery geometry, mass fraction, and homeostatic stress during PAH progression (Szafron et al. 2023). For a maladaptive alteration case in the pulmonary vascular tree associated with PAH, an additional variable representing an inflammatory stimulus is introduced and incorporated into the G&R function to simulate the progression of vascular maladaptation in PAH. However, a key limitation of these models is that they do not account for metabolic changes, such as the Warburg effect and mitochondrial dysfunction, which play a critical role in PAH progression.

Alongside structural remodeling, emerging evidences suggest that metabolic dysfunction is a pivotal role in the progression of PAH. One of the key drivers of this metabolic shift is the upregulation of hypoxia-inducible factor (HIF), which orchestrates cellular responses to chronic hypoxia by altering energy metabolism (Rai et al. 2008; Pullamsetti et al. 2020). Under hypoxic conditions, cells, including pulmonary artery smooth muscle cells (PASMCs), transition from mitochondrial oxidative phosphorylation to glycolysis, even in the presence of oxygen—known as the Warburg effect (Paulin and Michelakis 2014; Shi et al. 2020). This metabolic shift promotes increased glucose uptake and lactate production, which supports the hyperproliferation of PASMCs and endothelial cells, both of which contribute to vascular remodeling. Mitochondrial dysfunction in PAH further exacerbates this metabolic reprogramming by decreasing the activity of electron transport chain complexes I–III, impairing oxidative phosphorylation and other associated mechanisms, which leads to diminished ATP production (Shi et al. 2020; Riou et al. 2023). This shift to glycolysis is less energy-efficient, producing only 2 molecules of ATP per glucose molecule, compared to 36 ATP molecules generated by oxidative phosphorylation (Park et al. 2014). The metabolic alterations in PAH, therefore, play a critical role in driving both structural and functional changes in the pulmonary arteries, adding complexity to the task of accurately modeling PAH vasculature, as these metabolic shifts directly influence cellular proliferation, apoptosis resistance, and vascular remodeling (Xu et al. 2021).

Therefore, understanding of the relationship between metabolic process and pulmonary vascular wall remodeling is important for healthy and disease process. In fact, Murray’s law, which established a relationship between mechanical stress (e.g., wall shear stress) and metabolic energy consumption (termed cost) in the cardiovascular system, has been established (Taber 1998; Lindström et al. 2015). Particularly, in healthy pulmonary vasculature, computational frameworks have been established based on understanding of the metabolic energy optimization, in which Gharahi et al. (2023) presented a homeostatic optimization framework for a healthy pulmonary vasculature using an extension of Murray’s law, that optimizes geometry by incorporating metabolic energy costs and mechanical equilibrium constraints. Specifically, an iterative optimization, which integrating a metabolic cost function minimization, the stress equilibrium, and hemodynamics, has been performed at the slow timescale, while in the fast timescale, the pulsatile blood flow dimensional dynamics is described by a Womersley’s deformable wall analytical solution. The results of computational framework were compared with diverse literature data on morphometry, structure, mechanical behavior of pulmonary artery, and blood flow through the 1D vasculature. Nonetheless, the previous study was predicated on the assumption that the metabolic energy consumption of SMC and collagen fiber is unaltered on the vessel for a healthy subject. For PAH, the metabolic energy optimization with fixed metabolic energy consumption of SMC and collagen fiber is not an appealing choice, given the significance of metabolic alteration in PAH. However, quantitative data on metabolic energy consumption in vivo of the arterial vasculature are lacking, presenting a significant restriction on the use of the same optimization approach.

To address these limitations, we propose to develop a data-driven baseline model of PAH vasculature that integrates available geometric, pathological, and hemodynamic data from PAH patients. Baseline, in this context, refers to a state where the arterial geometry, hemodynamics, and wall properties represent those of a typical PAH patient. This baseline model will serve as a reference point from which changes induced by different PAH treatments can be measured. By establishing this baseline state, we aim to capture the specific characteristics of PAH-affected vasculature, allowing for the evaluation of treatment efficacy through the comparison of arterial and hemodynamic responses to interventions. This framework provides a critical foundation for assessing how different therapies modify vascular structure and function relative to this initial, disease-representative state.

The specific objectives of this work are twofold. Using the existing literature of experimental morphometric, pathological, and hemodynamic data, we first to construct a symmetrically bifurcating arterial tree, based on two assumptions: constant and varying wall constituents. This baseline model will allow us to simulate the mechanical behavior of PAH-affected arteries and study the impact of different hemodynamic conditions on vessel stiffness and stress. Second, while the metabolic costs of the healthy arterial wall have been extensively studied, there is a significant gap in our understanding of these costs in PAH. Therefore, we investigate to evaluate the metabolic costs for maintaining the arterial vasculature in PAH by testing two competing hypotheses on metabolic energy cost and discuss about plausible values for these energy expenditure in PAH.

## 2. Methods

Constructing a 1D-pulmonary vasculature and hemodynamic model for PAH is a multi-stage process aimed at capturing the complexity of arterial geometry, hemodynamics, and associated metabolic changes. In this study, we build on the 1D-pulmonary vascular model developed from Gharahi et al. (2023), which considered two cases for mass fraction assumptions: (1) a symmetric fractal tree with constant mass fractions of its wall constituents throughout the entire arterial network, and (2) a symmetric fractal tree with varying mass fractions across vessels of differing diameters. For the PAH model, we assume that the morphometry of arterial vasculature tree structure remained unchanged, except for modification in vessel thickness, internal diameter, and constituent mass fractions, with the values estimated from literatures. Once the PAH model is constructed, key arterial properties of vessels (e.g., stiffness, stress) and hemodynamic variables (e.g., blood flow, pressure, wall shear stress) are estimated throughout the vasculature tree as shown in **Figure 1**. In addition, metabolic energy costs are calculated under two conditions: (1) Unaltered metabolic energy cost of collagen-SMCs per unit volume, and (2) the same total metabolic cost of whole pulmonary vasculature as in the healthy vasculature.

**Figure 1.**
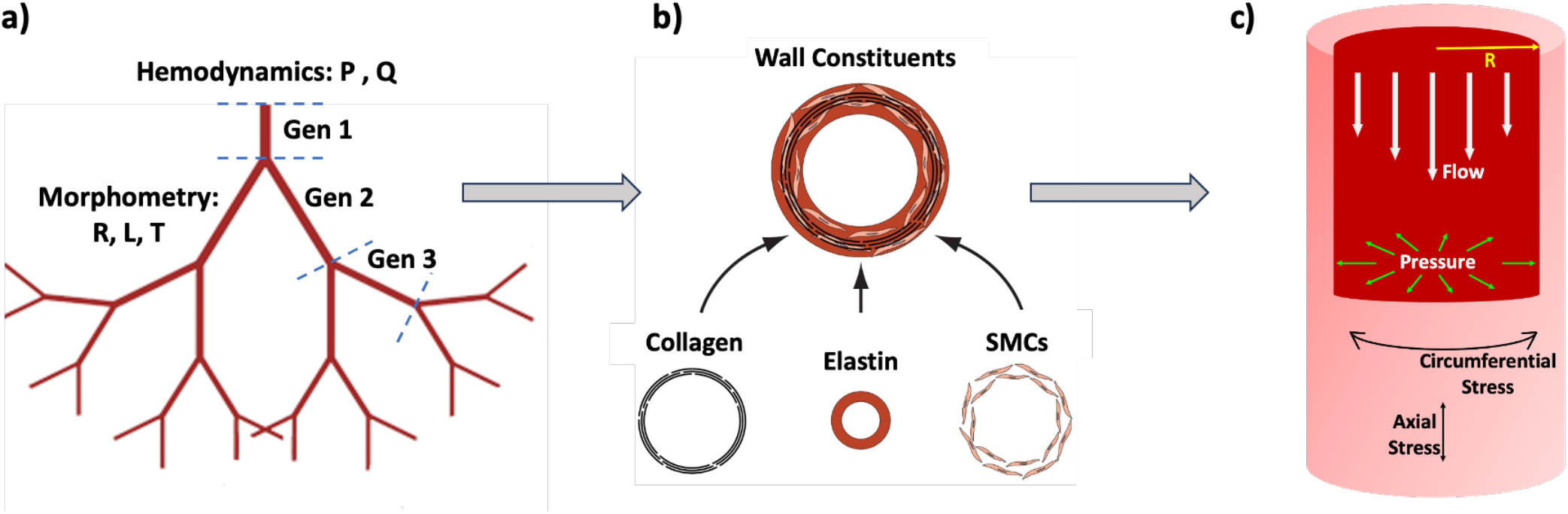
Schematic representation of establishment of baseline state for pulmonary arteries with PAH, a) morphometry (radius, length and thickness) and arterial tree hemodynamics (pressure and flow) are constructed, b) metabolic costs of wall constituents (elastin, collagen and smooth muscle cells) are estimated given with cases of vessel wall’s mass fractions, and c) arterial wall material and hemodynamic variables are calculated using a 1-D deformable hemodynamics model for the established PAH state.

### 2.1 Construction of Pulmonary Vasculature Model for PAH

#### 2.1.1 Morphometry of Arterial Tree

To establish the morphometry of the pulmonary arterial tree, we begin with healthy vasculature geometry and then, redefine wall thickness in accordance with the ratio of wall thickness to external diameters as reported in the literature data of PAH. We assume that the external diameter and length remain consistent between healthy and PAH vasculature, with increased wall thickness observed in PAH resulting in a reduced inner diameter and lumen. **Figure 2** illustrates the morphometric changes in the arterial wall from healthy and PAH-affected pulmonary arteries. In the healthy state (left), the arterial wall consists of thin intimal and medial layers, with a normal lumen size and regular organization of smooth muscle cells and ECM components. Conversely, the PAH state (right) exhibits significant remodeling, characterized by a thickened intimal layer due to endothelial cell proliferation and matrix deposition, as well as medial hypertrophy driven by smooth muscle cell proliferation and hyperplasia. The external diameter, 2 * *R*_*outer*_, remains largely unchanged, while the increased wall thickness, *T*_*PAH*_, reduces the lumen diameter, severely impairing blood flow. This structural remodeling underpins the elevated pulmonary vascular resistance and progressive functional decline observed in PAH.

**Figure 2.**
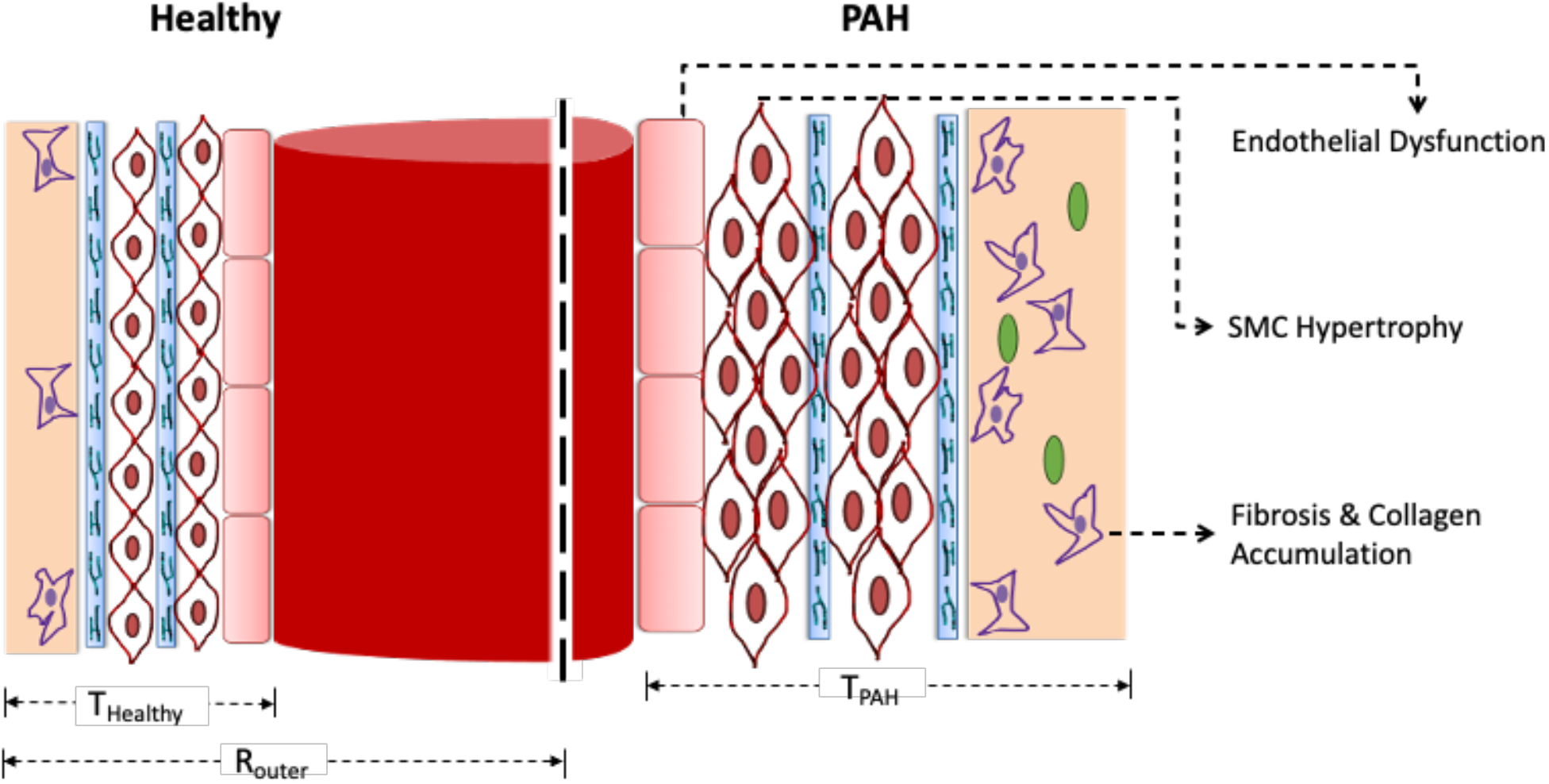
Schematic representation of the longitudinal cross-section of a healthy (left) and PAH (right) arterial wall.

The pulmonary vasculature model construction is briefly described. The exploration of vascular network architecture has long been inspired by Murray’s law, which suggests that vessels optimize their diameters to minimize energy dissipation. In our previous work (Gharahi et al. 2023) we extended Murray’s law to compute homeostatic radii for distal pulmonary vasculature, incorporating not only energy optimization for blood flow maintenance and viscosity dissipation-energy losses but also the energy for arterial wall maintenance, optimizing both radius and hemodynamics for healthy subjects.

In contrast to the Strahler ordering system (Jiang et al. 1994; Huang et al. 1996; Tamaddon et al. 2016), the morphometry of the tree is described by the generation number in this work, where the third bifurcated vessel from the main pulmonary artery is considered generation 1, and each bifurcation thereafter increments the generation number by 1, resulting in a total of 19 generations, as shown in **Figure 1(a)**. The initial stage focuses on accurately representing the geometric features of pulmonary arteries affected by PAH. The arterial geometry is, therefore, intended to replicate the pathological changes that have been documented in the literature, thereby providing a foundation for subsequent modeling stages (Santos et al. 2002).

Starting with the previous work (Gharahi et al. 2023), we maintained the outer diameter similar to that of healthy arteries across different vessel generations. This approach is based on the complexity of the relationship between generation number and diameter measurement in histology observations from human-tissue data. Although real pulmonary arteries have asymmetric bifurcations, we presume that the outer diameter of pulmonary arteries remains the same as that of healthy vasculature, in which this assumption provides a logical and simplified starting point for modeling. From this, the wall thickness and inner diameter for PAH arteries are then estimated using an empirical relation that was fitted to histologically measured data for PAH vasculature at autopsy (Rol et al. 2017), which provided detailed measurements of the inner and outer diameters of pulmonary arteries in six PAH patients. By fitting the dataset with a linear regression, an empirical relation between the inner and outer diameters is derived, reflecting the average inner-to-outer diameter ratio across these six patients. The individual linear fits for each patient showed variability, with slopes ranging from 0.33 to 0.84, highlighting the heterogeneity of vascular narrowing in PAH. Averaging these slopes led to the inner-outer diameter relation *D*_*i*_ = 0.66 * *D*_*o*_, which allows for the estimation of the inner diameter (*D*_*i*_) and wall thickness (H) in PAH vasculature:

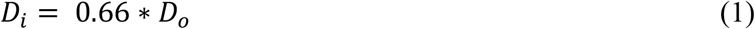

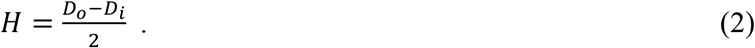

This approach enables a more realistic representation of PAH progression by maintaining structural consistency with healthy vasculature while incorporating the specific changes in wall thickness and inner diameter observed in PAH, as informed by the histological data.

The length of arterial vessel is related to the radius using the relation given in (Olufsen et al. 2012) which is an empirical relation as shown below utilizing the morphometric data from (Huang et al. 1996), where the length of arterial vessel (*L*) and the vessel inner radius (*R*) are measured in *mm*.

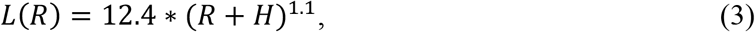

where *H* is the vessel wall thickness. The data used in this reference primarily focuses on morphometric characteristics in healthy subjects, but the relation has been extended for use in diseased conditions like PAH. This length-radius relation allows us to model the geometry of the pulmonary arteries in both healthy and diseased states, providing insights into how vascular remodeling in PAH affects the overall geometry of the arterial network.

Using the above relations, morphometry of the entire arterial tree (radius, thickness and length) is defined for the entire PAH vasculature, where radius, thickness and length are plotted with respect to the external diameter and arterial generation numbers, shown in **Figure 3**.

**Figure 3.**
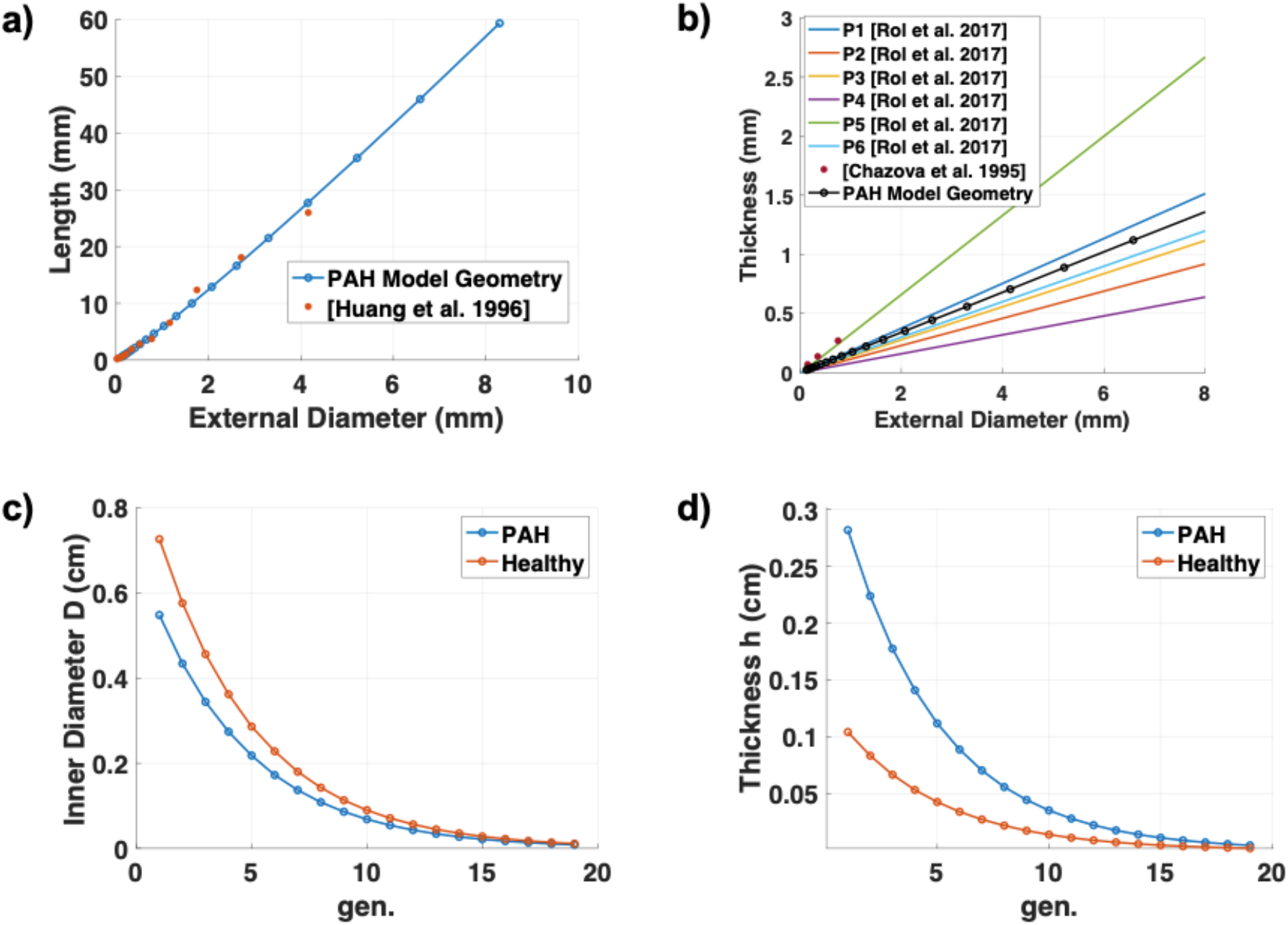
Morphometry of PAH arterial tree compared to literature and previously established healthy vasculature. a) Length of the arterial tree for different diameters as compared with the histological data from (Huang et al. 1996), b) thickness of vessel wall at different external diameters fitted in the model as compared with the PAH data from different patients (Chazova et al. 1995; Rol et al. 2017), c) inner diameter, and d) thickness of the arterial tree across different generations are plotted and compared to homeostatic healthy vasculature as described in (Gharahi et al. 2023).

#### 2.1.2 Mass Fractions of Arterial Wall

The two cases of baseline states in this current study are: 1) symmetric arterial tree with constant mass fraction 2) symmetric arterial tree with varying mass fractions through the vasculature. The total mass per unit areal cross section for each generation of the arterial tree is related to thickness as given below

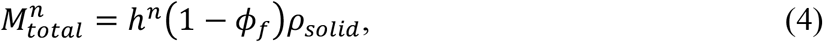

where *h*^*n*^ is the total thickness of *n*th generation arterial wall, *ϕ*_*f*_ is the volume fraction of the interstitial fluid, usually taken as 0.7, and *ρ*_*solid*_ is the density of the dried arterial wall, which is considered as 1060 kg/m^3^. In our model, vessel wall is assumed to be made up of only 3 constituents-elastin, smooth muscle cells and collagen fibers, where the total mass per unit areal area is divided by

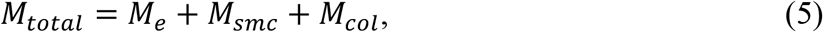

where *M*_*e*_, *M*_*smc*_, and *M*_*col*_ represent the mass per unit area cross section of elastin, smooth muscle cells and collagen fibers, respectively.

For the healthy pulmonary vasculature, Gharahi et al. (2023) constructed the model with 2 cases on the constituents of arterial wall, made estimation/assumptions based on literatures. For the first case of assuming a constant mass fraction, intimal layer thickness and its constituents are neglected as it contributes to <15% of total thickness (Chazova et al. 1995) and adventitial layer was assumed to be comprised of 95% of collagen and 5% of elastin, neglecting other constituents such as fibroblasts and endothelial layer for ease of modelling. Combined with these assumptions and reported mass fractions of the constituents in the medial layer as reported by Mackay and colleagues (Mackay’ et al. 1978), the constant mass fractions for a healthy vasculature as shown in **Figure 4(a)** are defined.

**Figure 4.**
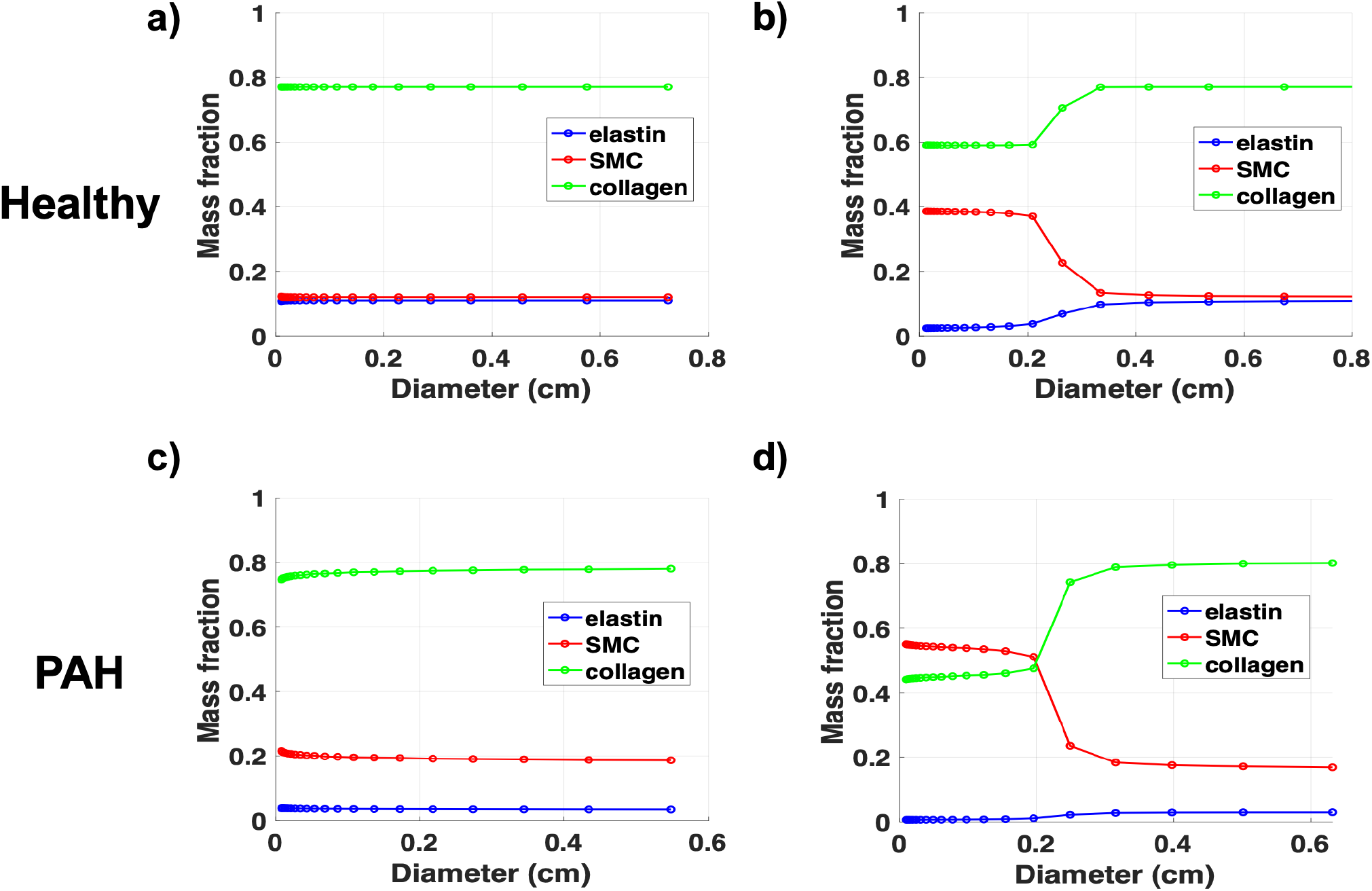
Mass fractions of constant and variable mass fractions across different diameters for (a) & (b) healthy and (c) & (d) PAH.

However, various pulmonary artery pathology studies (Elliott and Reid 1965; Chazova et al. 1995; Townsley 2012) have shown that the mass fractions of wall constituents vary from the main pulmonary arteries to capillary arteries, transitioning from elastic to transitional, muscular, partially muscular, and non-muscular arteries. This study however excludes arterial vessels with smaller diameter of less than 0.01 *cm*, instead focusing on the region of large and small vasculature in the transition between elastic and muscular arteries as highlighted in **Figure 4(b)**. To establish the mass fractions of a PAH arterial tree, the following assumptions are made: 1) mass of elastin (M_e_) is assumed to be same as a healthy arterial wall for the comparable generation, 2) change in medial layer thickness is contributed fully due to increase in mass of smooth muscle cells (M_smc_). The second assumption is motivated by the studies (Tobal et al. 2021). Along with the above assumptions and histological data in change in wall thickness for PAH (Chazova et al. 1995), mass fractions for a PAH arterial wall are established for both the cases as shown in **Figure 4(c)** and **4(d)**.

#### 2.1.3 Hemodynamics with PAH

In parallel with the geometric and mass fraction model construction, the model incorporates the hemodynamic aspects specific to PAH as described in (Lankhaar et al. 2006; Zambrano et al. 2018). Hemodynamic simulations within the computational model aim to replicate the dynamic conditions within PAH-affected pulmonary arteries. Along with slow-time hemodynamics (minutes), pulsatile hemodynamics are evaluated using the fluid-solid-growth (FSG) framework and Womersley’s solution (Figueroa et al. 2009; Filonova et al. 2020; Gharahi et al. 2023) as briefly described in **Supplementary Materials**. From the PAH hemodynamics data, the inlet flow and pressure are described at the boundaries, which is used to solve the hemodynamics conserving flow and maintaining constant pressure at the bifurcation as shown in **Figure 5(c)** below.

**Figure 5.**
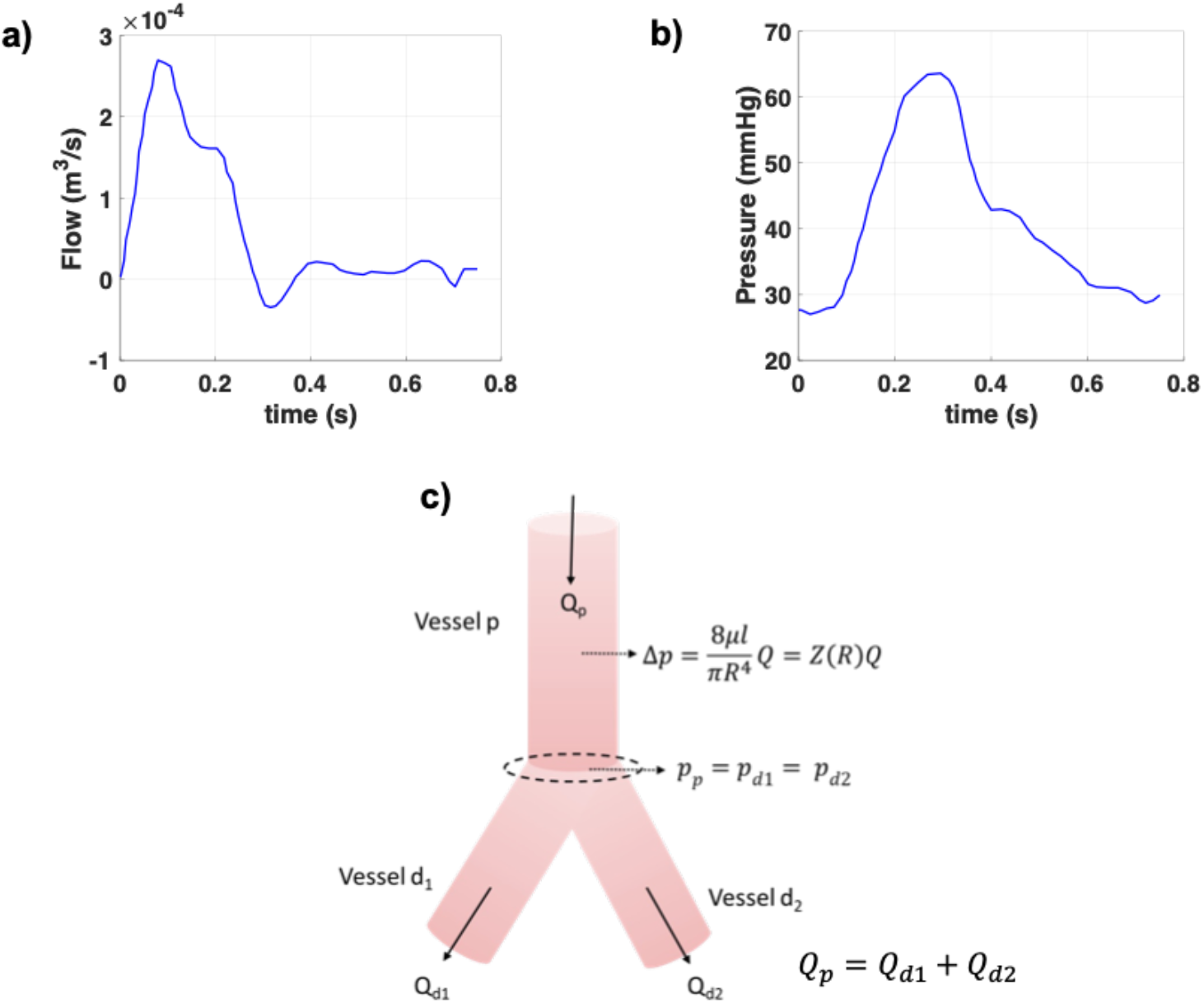
a) Flow profile and b) pressure profile of a PAH patient as reported in (Lankhaar et al. 2006), c) hemodynamics in an arterial vessel bifurcation.

Using the pressure and flow profiles of the main pulmonary artery (MPA) from hemodynamic measurements of a PAH patient (Lankhaar et al. 2006), as shown in **Figure 5 (a)** and **(b)**, we extend these to the 1st generation (third bifurcation from the MPA) in our model with the following assumptions: (1) the pressure drop is negligible across the first three bifurcations, so the input pressure for the 1st generation is the mean pressure from the MPA profile; and (2) flow is halved at each bifurcation in a symmetric tree, making the input flow for the 1st generation 1/8th of the MPA mean flow. After defining the input flow and pressure at the 1st generation of the arterial tree, the pressure, flow, and resistance across all subsequent generations can be evaluated using the following relations:

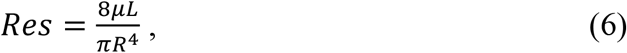

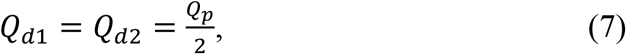

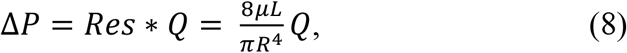

where *Res* is the resistance, *R* is the radius of the blood vessel, *Q* is the blood flow, and *μ* is the dynamic viscosity of blood taken as 0.0035 Pa sec *μ*.

### 2.2 Metabolic Cost of PAH Arterial Tree

This section explains how metabolic cost and energy are calculated for given the arterial geometry, hemodynamics, and mass fractions, in contrast to the previous study (Gharahi et al. 2023), where the metabolic costs were defined from experimental data and the geometry was computed using the metabolic energy cost optimization method. Based on (Gharahi et al. 2023), the total metabolic energy cost (*C*) of the pulmonary vasculature per unit length for an individual blood vessel can be defined and calculated as

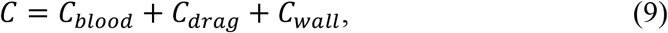

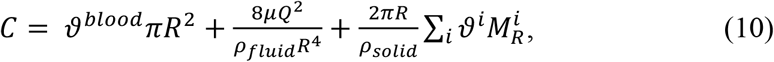

where *ρ*_*fluid*_ is the blood density which is 1060 kg/m^3^, *ϑ*^*blood*^ is the metabolic energy consumption of blood supply which is 51.7 W/m^3^ (Liu and Kassab 2007), *ϑ*^*i*^ is the metabolic energy consumption of vessel wall constituent *i*, which is considered to be 0 W/m^3^ for elastin, 1500 *W/m*^*3*^ for collagen and smooth muscle cells maintenance, and 0.00872 s^-1^ for active tension(Liu and Kassab 2007; Paulin and Michelakis 2014). Using the above relations, we calculate the total metabolic energy cost based on two competing hypotheses: 1) calculating the energy cost *C* per unit length while keeping the metabolic energy consumption of the constituents *ϑ*^*i*^ for unit volume the same as healthy arterial wall and 2) calculating the energy consumption of the constituents *ϑ*^*i*^ for each arterial generation by adjusting the energy consumption *ϑ*^*i*^, while the total energy cost *C* per unit length is assumed to be same as a healthy vessel.

## 3. Results

Utilizing the presented data-based computational model, key mechanical parameters and hemodynamic variables are predicted based on two scenarios with symmetric constant and variable mass fractions and compared for healthy and PAH-affected pulmonary arteries. This comparative analysis highlights the deviations in mass distribution, emphasizing the impact of PAH-induced geometric and hemodynamic changes. Understanding the differences in mass fractions provides valuable insights into the disease progression and aids in the identification of key parameters influencing PAH.

### 3.1 Arterial Wall Mechanics

The results of computational analysis of vascular arterial mechanical and slow-time hemodynamics (mean flow and pressure) parameters are presented for arterial stiffness, wall shear stress, and linearized modulus (Baek et al. 2007; Gharahi et al. 2023). The comparison between the stiffness of healthy arteries and PAH arteries (variable stiffness case) shows a drastic increase in PAH. In **Figure 6(a)** and **(c)**, the stiffness for the healthy case remains around 7-10 kPa across generations, aligning with previous experimental values (Krenz and Dawson 2003).

**Figure 6.**
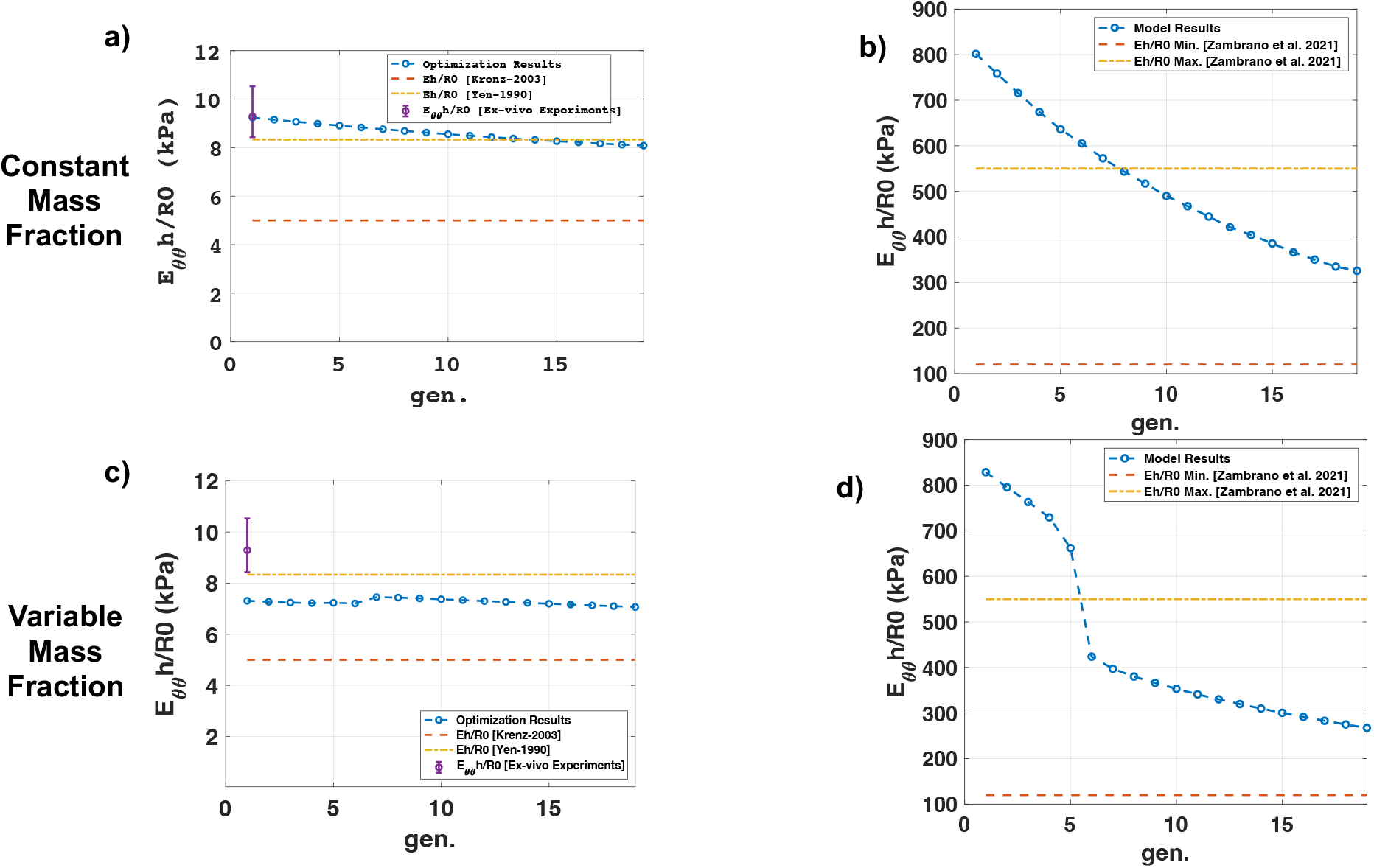
Arterial wall stiffness across different generations for healthy and PAH case with constant and variable mass fractions compared with the literature data. Ex-vivo experiments were the values computed from (Yen et al. 1990; Krenz and Dawson 2003; Zambrano et al. 2021; Wang et al. 2021)

In contrast, in the PAH case **Figure 6 (b)** and **(d)**, the stiffness is significantly higher, starting around 800 kPa and decreasing to about 300 kPa over the generations, which is approximately 10 times higher than in the healthy condition. This stark increase illustrates the heightened arterial stiffness in PAH due to vascular remodeling and loss of compliance.

In **Figure 7**, the wall shear stress (WSS) *τ* is plotted against vessel outer diameter (D) for both healthy and PAH conditions. In **Figure 7(a)** (healthy, constant mass fraction), shear stress starts at approximately 1.2 Pa for larger diameters and increases to around 2.0 Pa for smaller diameters. Similarly, in **Figure 7(c)** (healthy, variable mass fraction), the trend is consistent with a slight rise in shear stress at smaller diameters.

**Figure 7.**
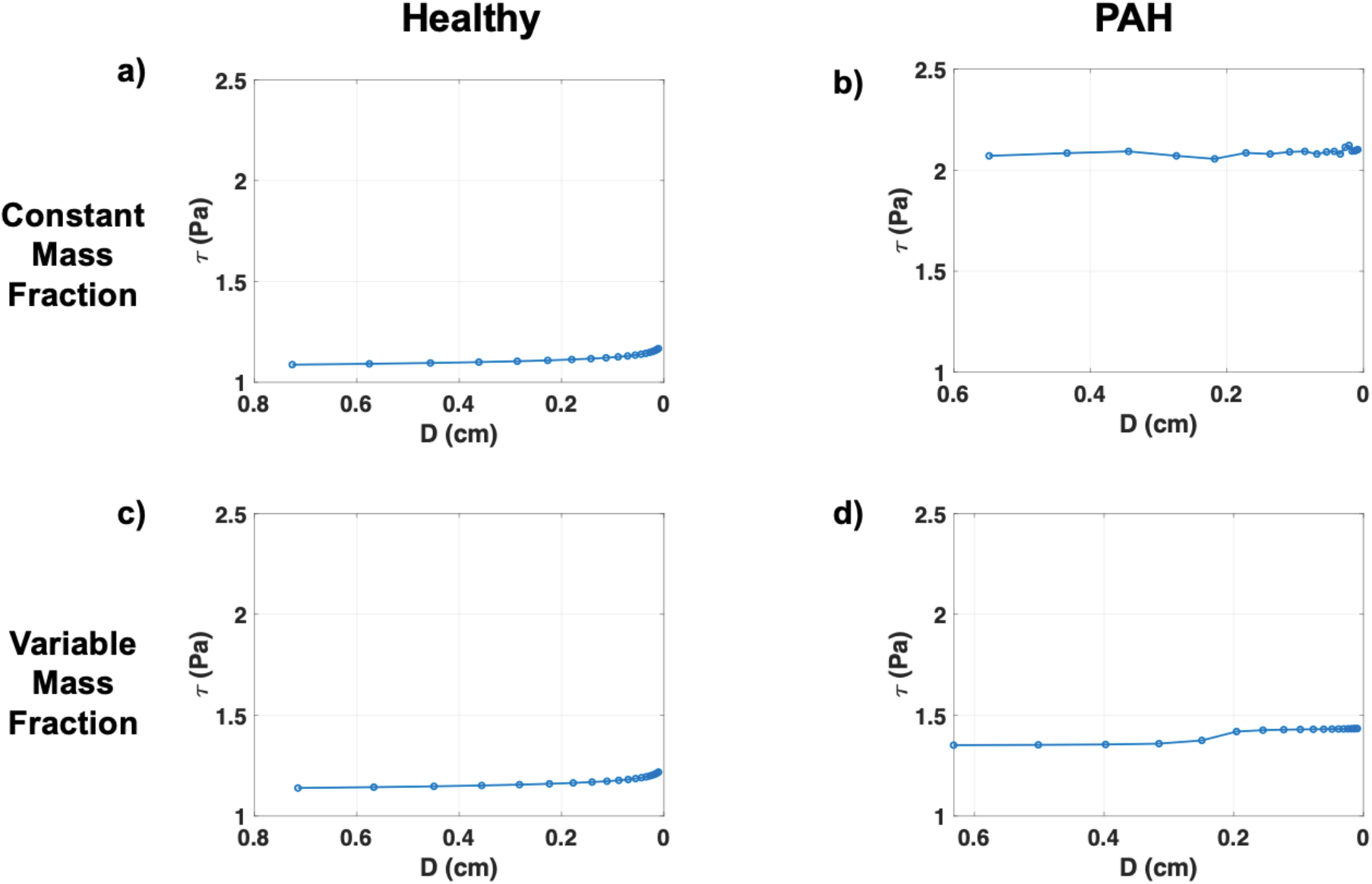
Shear stress across different generations for healthy and PAH case with constant and variable mass fractions.

For PAH, in **Figure 7(b)** (constant mass fraction), shear stress is notably higher and more stable, starting around 2.0 Pa across all diameters, while in Figure. 3.10 d) (variable mass fraction), it shows a more gradual increase from about 1.5 Pa to 1.8 Pa as the diameter decreases. These results indicate that PAH leads to higher and more uniform shear stress across generations compared to healthy conditions. For healthy pulmonary arteries, the results in **Figure 7(a)** and **(c)** indicate that WSS increases as vessel diameter decreases, a trend consistent with normal hemodynamic behavior and flow distribution in arterial trees. The presence of constant or variable mass fractions does not significantly alter this gradient under healthy conditions. Such findings align with prior studies emphasizing the physiological role of WSS in maintaining endothelial homeostasis and nitric oxide (NO) production, which collectively ensure vascular stability. In PAH conditions, as shown in **Figure 7(b)** and **(d)**, elevated pressures in the proximal pulmonary arteries result in an overall higher baseline WSS compared to healthy arteries.

In **Figure 8**, the graphs compare the (linearized) Young’s modulus across generations for both healthy and PAH conditions. In **Figure 8(a)** (for a healthy subject and constant mass fraction), the modulus starts around 50 kPa and gradually decreases to about 35 kPa by the later generations. In **Figure 8(b)** (PAH, constant mass fraction), the modulus is significantly higher, starting at approximately 700 kPa and declining steadily. In **Figure 8(c)** and **(d)** (PAH with variable mass fraction), the modulus exhibits a distinct drop between generations 5 and 6, highlighting the drastic stiffening and mechanical alterations in the PAH vasculature.

**Figure 8.**
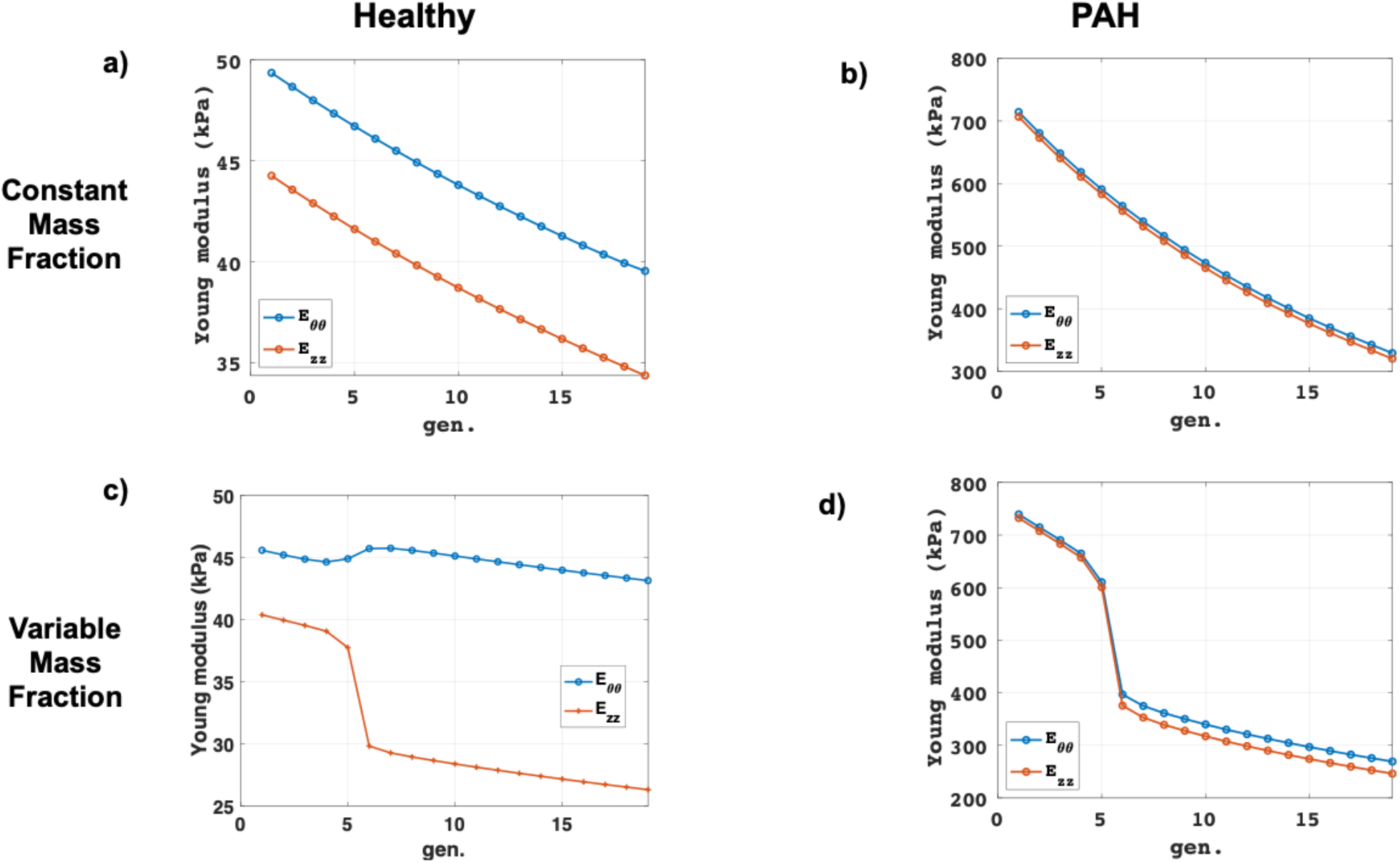
Youngs modulus across different generations for healthy and PAH case with constant and variable mass fractions.

### 3.2 Pulsatile Hemodynamics Using Womersley’s Solution: Healthy vs. PAH

Pulsatile hemodynamics results for the healthy and PAH vasculature for symmetric arterial tree with constant mass fractions. The graphs in **Figure 9** displays both the flow and pressure profiles along generations of the pulmonary arterial tree under healthy and PAH conditions. For healthy arteries, the flow gradually reduces from the proximal to distal generations, showing a clear pattern of decreasing amplitude, while the pressure also declines steadily along the arterial network, shown in **Figure 9(a)**. In contrast, in the PAH condition, flow exhibits greater irregularity characterized by a significant reversed flow at *t*=~0.5 *sec* and reduced amplitude in later generations. Pressure remains significantly elevated and less dynamic throughout the generations in PAH, indicating increased vascular stiffness and reduced compliance, which is the key characteristics of PAH-induced remodeling.

**Figure 9.**
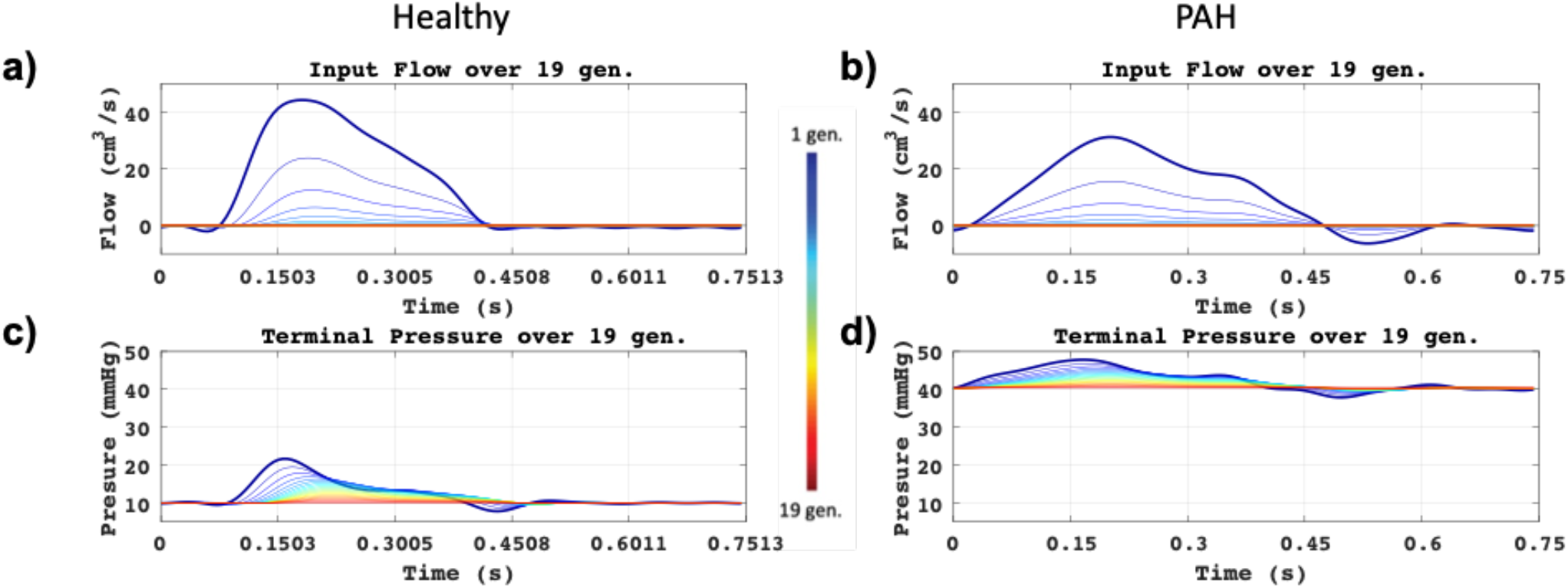
Pulsatile hemodynamics of healthy and PAH vasculature a) & b) Flow profile and c) & d) pressure profile.

**Figure 10.**
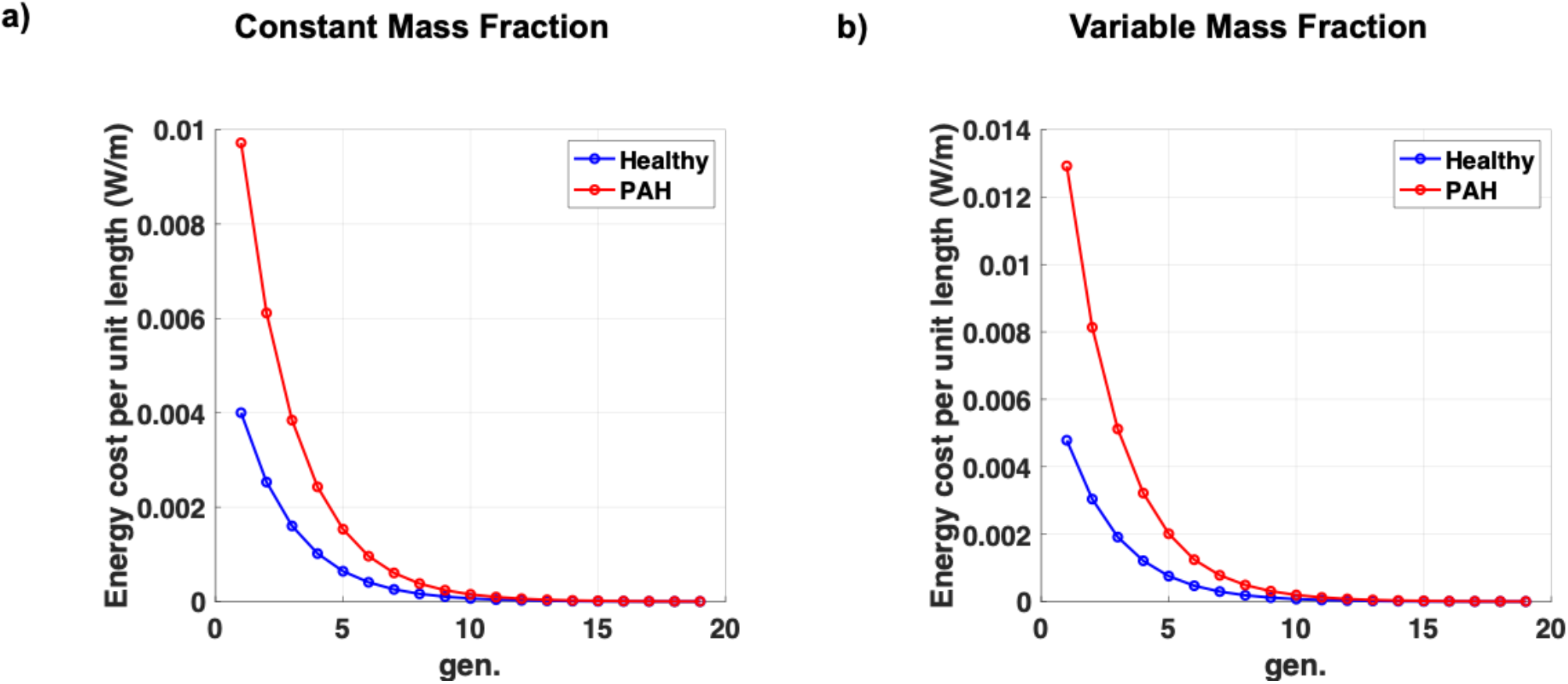
Energy cost per unit length across different generations of arterial tree for healthy and PAH vasculature for a) constant and b) variable mass fractions.

### 3.3 Metabolic Energy Cost of the Arterial Segment

The metabolic energy cost for the arterial segment is computed for the two cases of constituent’s mass fraction (constant vs variable) for both healthy and PAH vasculature, assuming the constant metabolic energy cost of arterial constituents per unit volume, plotted in **Figure 10(a)** and **(b)**. For the constant constituent’s mass fraction, where the metabolic energy consumption of the constituents *ϑ*^*i*^, same as healthy arterial wall, the metabolic energy cost per unit length is plotted against the generation numbers for the PAH and healthy subject. It is computed at approximately 0.01 W/m for unit length in the first generation and gradually decreases across the subsequent generations, while for the healthy vasculature, it begins at around 0.004 W/m in the first generation. The computed results show that the energy cost in the PAH case is more than twice that of the healthy case, especially in the initial generations. The computational results from the first hypothesis leads that the pulmonary vasculature consumes a twice amount of energy, which may be unreasonable due to the less energy-efficient metabolic process associated with PAH (Peng et al. 2016).

In contrast, for the second hypothesis as shown in **Figure 11**, the metabolic cost is evaluated assuming that the energy cost per unit length is same as that of healthy. In the case of constant mass fraction, the metabolic consumption increases linearly across generations, starting at approximately 500 W/m^3^ and rising to about 625 W/m^3^. For arterial tree with variable mass fraction, the metabolic consumption remains relatively stable across generations, fluctuating around 450-500 W/m^3^. This indicates that by reducing the metabolic consumption, the overall energy cost expenditure becomes comparable to that of healthy arteries.

**Figure 11.**
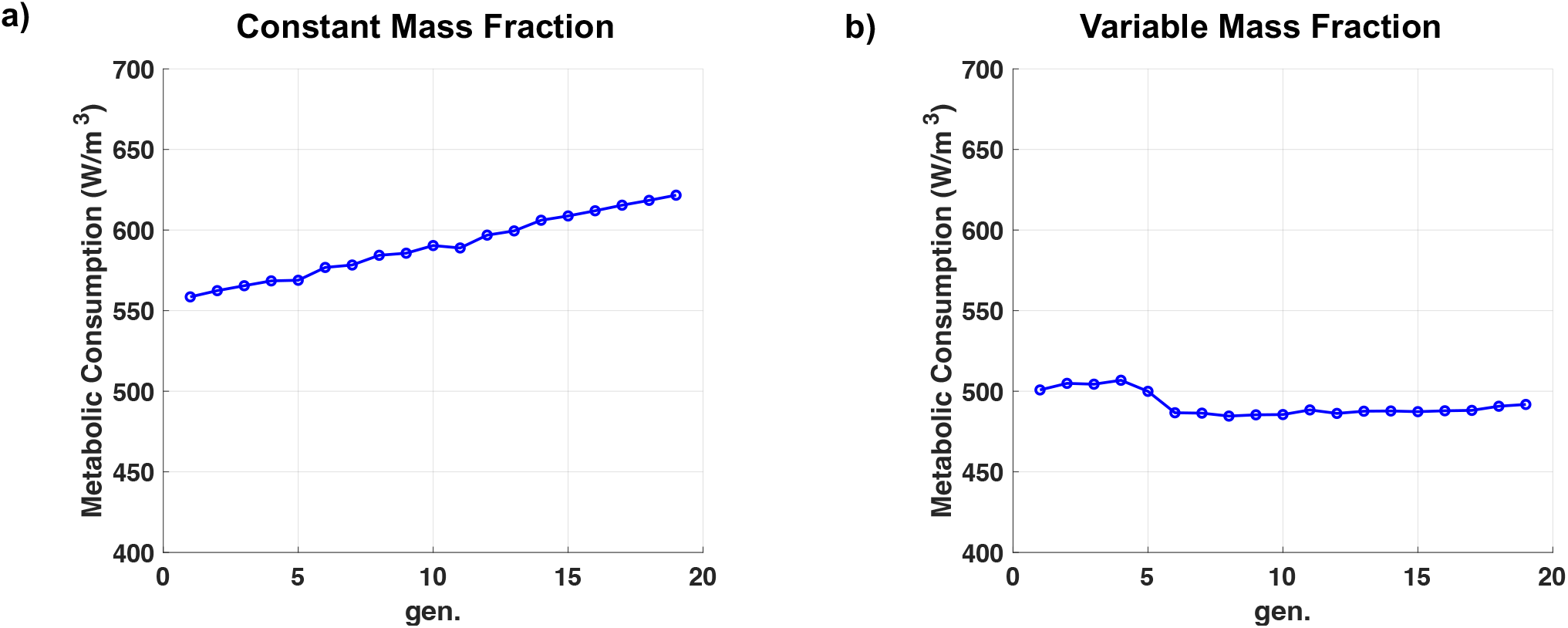
Metabolic consumption of arterial wall across different generations of arterial tree for healthy and PAH vasculature for a) constant and b) variable mass fractions.

## 4. Discussion

A computational vascular model should accurately depict physiologically realistic tree morphometry, as it affects vascular resistance and impacts long-term vascular adaptation and hemodynamics during the cardiac cycle. However, creating a complete and physiologically precise morphometry of the distal pulmonary arterial tree poses a difficulty due to limitations in data acquisition and the complexity of modeling (Burrowes et al. 2005). Inspired by Murray’s law, which optimizes the metabolic energy cost in biological systems and serves for identifying bifurcation patterns (Murray 1926), several approaches have been proposed to characterize vascular network architecture. These include fractal-based morphometry (Zamir 1999; Kamiya and Takahashi 2007; Ionescu et al. 2009) and multiscale hemodynamic and fluid-solid interaction (FSI) analyses (Olufsen et al. 2000; Mittal et al. 2005; Van De Vosse and Stergiopulos 2011; Qureshi et al. 2014). Nevertheless, these studies often rely on simplified vessel wall representations that do not account for the contributions of individual structural constituents—an essential factor in understanding structural changes associated with PAH. To address this limitation, Gharahi et al. (Gharahi et al. 2020, 2023) developed computational models that enable long-term hemodynamic and fast hemodynamics with FSI analysis, while integrating physiologically realistic tree morphometry and structurally motivated vessel wall models that estimate the level of healthy homeostatic stresses and hemodynamic variables.

Building upon Gharahi’s framework, this study presented a baseline model of PAH vasculature, defining key geometric, mechanical, and hemodynamic variables while estimating the metabolic characteristics of affected arteries. It also compares critical parameters—such as arterial stiffness, wall shear stress, and energy costs—between healthy and PAH-affected arteries. This hybrid (data-driven and computational) baseline model accurately reflects key morphometric changes observed during disease progression. By incorporating variations in wall thickness, lumen narrowing, and hemodynamic parameters, the model enables a detailed analysis of disease-specific mechanical and structural properties of the pulmonary vasculature. Establishing baselines for both healthy and diseased pulmonary arteries reveal significant differences in geometric, mechanical, and hemodynamic variables.

The first key finding of computational results is that arterial stiffness in PAH-affected vessels is nearly ten times higher than in healthy arteries, indicating a severe loss of elasticity. The computational results align with these observations, such as reported the structural stiffness (*E*_*θθ*_*h/R*_0_) 7.1 to 9.3 kPa for healthy subjects (Yen et al. 1990; Krenz and Dawson 2003) and in the range of 280 to 840 kPa in PAH patients (Zambrano et al. 2021) shown in **Figure 6**, demonstrating a comparable magnitude for the Young’s modulus across arterial generations. The computed axial and circumferential moduli exhibit trends consistent with experimental data, showing increased stiffness in proximal vessels, which decreases progressively along the distal pulmonary arterial tree. Other studies on pulmonary arteries in PAH-afflicted calves reported that the Young’s modulus under PAH conditions is approximately three to four times higher than in normal arteries, with values ranging from 100 to 200 kPa—substantially exceeding those in healthy pulmonary arteries with the values ranging from 50 to 100 kPa (Lammers et al. 2008). These findings align with calf models of PAH, where increased vascular stiffness is a hallmark of disease-induced remodeling. For example, mean elastic modulus values in PAH conditions have been reported as 192 ± 56.7 kPa, compared to 97 ± 30.1 kPa in healthy calf controls (Hunter et al. 2010a).

Secondly, a difference in WSS distribution is observed between healthy and PAH-affected arteries. WSS plays a crucial role in normal physiological mechano-sensing and is linked to pathological signaling and disease progression (Wang and Valdez-Jasso 2021; Allen et al. 2023). Under PAH conditions, as shown in **Figure 7(b) and (d)**, elevated pressures in the proximal pulmonary arteries lead to an overall increase in baseline WSS compared to healthy arteries. The increase in wall shear stress (WSS) observed in our computational simulations is consistent with computationally estimated results by (Yang et al. 2019; Szafron et al. 2024), who reported elevated WSS in pathological vascular remodeling, further supporting the concept that mechanical forces significantly influence vascular adaptation in PAH. In smaller, distal vessels, heightened WSS may exacerbate endothelial injury and trigger inflammatory signaling, promoting pathological changes such as intimal hyperplasia and medial thickening (Humbert et al. 2019). In contrast, larger, proximal arteries in PAH cases experience reduced WSS, contributing to endothelial dysfunction through impaired NO production and upregulation of pro-inflammatory mediators. This dual pattern, increased WSS in distal arteries and decreased WSS in proximal arteries, highlights the complexity of hemodynamic forces driving disease progression (Tang et al. 2012). The variability of WSS across the pulmonary arterial tree, as depicted in the computational model, aligns with advanced imaging studies such as those utilizing four-dimensional flow MRI. From a therapeutic perspective, normalizing WSS presents a promising strategy to mitigate PAH progression. Pharmacological interventions such as PDE5 inhibitors and prostacyclin analogs (Tettey et al. 2021), which reduce pulmonary arterial pressures, may indirectly restore WSS to more physiological levels, thereby improving endothelial function. Emerging treatments like Sotatercept, which target structural remodeling (Doggrell 2023), may further enhance the normalization of WSS.

Lastly, the computation of metabolic energy cost *C* per unit length derived from the assumption that the metabolic consumption *ϑ* by the constituents is same as that of healthy arterial wall yields double the total metabolic energy cost for the first hypothesis, whereas the metabolic energy consumption *ϑ* of vessel wall per unit volume significantly decreases when the total energy cost *C* per unit length is presumed to be equivalent to that of a healthy vessel. Maintaining the same metabolic consumption *ϑ* per unit volume is likely to result in excessive metabolic energy cost expenditure, imposing significant stress on patients’ bodies; thus, reducing metabolic consumption of vessel wall per volume may be advantageous from the energy cost expenditure perspective. Particularly to the case of arterial tree with variable mass fraction, the results of metabolic consumption (per unit length) remain relatively stable across generations, fluctuating around 450-500 W/m^3^, shown in **Figure 11(b)**. This indicates that by reducing the metabolic consumption of vessel wall, the overall energy cost expenditure becomes comparable to that of healthy arteries.

While the Warburg effect was primary studied in cancer research (Benny et al. 2020), by the early 2010s, the heightened activity of pyruvate dehydrogenase kinase (PDK) was implicated in cardiac hypertrophy (Piao et al. 2010), potentially leading to heart failure, broadening the scope of the Warburg phenomenon to other conditions (Dabral et al. 2019; Huang et al. 2022). The role of hypoxia signaling gained prominence in both non-malignant and malignant hyperproliferative vascular diseases like pulmonary hypertension, atherosclerosis, and vascular restenosis. It has been suggested that metabolism is modified in stressed conditions such as hypoxia, oxidative stress, and inflammation, being altered from usual mitochondrial oxidative phosphorylation to lower-efficient glycolysis (Piao et al. 2010; Dabral et al. 2019; Benny et al. 2020; Huang et al. 2022). Especially, it has been suggested that a key feature of PAH is the reduced contractility of PASMCs under hypoxic conditions, largely driven by mitochondrial dysfunction. In PASMCs isolated from PAH patients, the expression of mitochondrial ETC components I–V is significantly reduced, leading to decreased mitochondrial oxygen consumption (Shi et al. 2020). In animal models, the activity of mitochondrial respiratory chain complexes I–III is reduced by approximately 50%, correlating with a similar reduction in overall mitochondrial oxygen consumption (Shi et al. 2020). This can be corroborated with the reduction in metabolic consumption as shown in **Figure 11**. This dysfunction in mitochondrial oxidative phosphorylation directly impacts PASMC contractility by limiting ATP production, which is critical for muscle function. Reduced energy availability under hypoxia leads to diminished contractility and promotes the onset of vascular remodeling characteristics of PAH.

During the early stages of PAH, PASMCs exposed to chronic hemodynamic stress exhibit further declines in mitochondrial oxidation and endothelial cell activity, exacerbating the disease’s pathophysiology. The energy deficit caused by mitochondrial impairment not only weakens contractility but also drives PASMC proliferation, contributing to arterial thickening and increased pulmonary vascular resistance (Marshall et al. 2018). Hypoxia plays a pivotal role in regulating these metabolic shifts, pushing PASMCs toward a glycolytic phenotype that further disrupts their normal contractile function (Taggart and Wray 1998). Mitochondrial dysfunction in PAH also involves hyperpolarized mitochondria, which suppresses apoptosis and promotes cell proliferation. This is accompanied by reduced production of mitochondrial-derived reactive oxygen species (mROS) and activation of hypoxia-inducible factor (HIF)-1α, which stimulates glycolysis (Ghofrani et al. 2011; Xu et al. 2021).

Beyond glucose metabolism, the metabolic alterations in PAH extend to lipid and amino acid metabolism. Non-oxidized sugars, lipids, and amino acids are redirected toward biosynthetic pathways, which support cellular proliferation and survival (Paulin and Michelakis 2014); This altered metabolic state contributes to the resistance to apoptosis, a hallmark of PAH, and promotes the extensive vascular remodeling observed in the disease. Recent studies also suggest that these metabolic changes activate epigenetic mechanisms and inflammation via mitochondria-based inflammasomes, linking metabolic dysfunction to the progression of vascular remodeling (Xu et al. 2021).

The significance of this model lies in its potential as a platform for future integration of pharmacological pathways and treatment simulations. While this study focuses on establishing the baseline network, the framework is inherently designed to accommodate additional pathways, such as the nitric oxide-cGMP-PKG signaling cascade (Mullagura et al. 2021), enabling the exploration of therapeutic interventions and patient-specific treatment predictions. The ability to extend this baseline into a predictive model for treatment response offers a transformative tool for advancing the understanding of PAH progression and its management. By providing a detailed, data-driven representation of the diseased arterial network, this study bridges an important gap between theoretical modeling and experimental observations. This alignment with existing experimental and clinical data enhances the reliability of the framework and underscores its potential to serve as a reference for further studies on vascular remodeling. Additionally, this baseline model is significant for its versatility. Beyond its utility in PAH research, it serves as a foundation for broader applications in vascular diseases involving remodeling and altered hemodynamics. Its modular structure makes it adaptable for investigating interactions between biomechanics and biochemical pathways, positioning it as a critical resource for interdisciplinary studies. Overall, the established model sets the stage for integrating biochemical pathways and pharmacological simulations, paving the way for the development of predictive tools to optimize treatment strategies for PAH.

While this study presents a significant step forward in modeling the baseline PAH arterial network, several limitations must be acknowledged. One limitation arises from the assumptions and idealizations made during the modeling process. For example, the geometrical and structural parameters used to represent the pulmonary arterial network are derived from limited datasets and may not capture the full variability observed across different patients and disease stages. The simplifications, while necessary to build a tractable model, may exclude some intricate features of vascular remodeling, such as localized heterogeneity in wall thickening or regional variations in stiffness (Rol et al. 2017). Another limitation is the gap in experimental data on metabolic costs specific to PAH. While the model incorporates hemodynamic and structural changes, the lack of comprehensive data on the metabolic energy consumption of PASMCs in diseased states prevents a deeper integration of energy dynamics into the framework. This gap restricts the ability to fully capture the energetic shifts, such as those driven by the Warburg effect, in the pathophysiology of PAH. Furthermore, the model assumes a standardized disease state, which poses challenges when addressing the heterogeneity of disease progression. The progression and severity of PAH vary significantly among patients, influenced by factors such as genetics, comorbidities, and environmental exposures (Talwar et al. 2017). This variability makes it difficult to develop a single model that encompasses all degrees of disease progression. As such, while this model serves as a robust baseline, its predictive capacity may be limited for extreme or atypical cases. Despite these limitations, the model provides a valuable foundation for advancing PAH research. Its modular design would allow for the integration of additional data and pathways as they become available, enabling refinement and expansion over time (Savale et al. 2018). While no model can fully capture the complexity of PAH, this study marks an important step forward, providing a framework that can guide further research and facilitate the development of targeted therapies.

In conclusion, the construction of a comprehensive baseline model for PAH involves a meticulous process spanning arterial geometry, hemodynamics, and metabolic considerations. The three stages—geometry construction, pathological estimation, and metabolic costs analysis—ensure a robust representation of PAH arterial mechanics. The findings highlight the significant alterations in mechanical properties, particularly the increased stiffness and energy expenditure in PAH vasculature. These insights are crucial for advancing our understanding of PAH pathophysiology and for developing future treatment strategies. The model offers potential for simulating patient-specific arterial responses to treatments, contributing to a more effective clinical approach in managing PAH.

## Appendix

### A. Womersley Theory for Deformable Walls

Womersley’s theory, initially developed for rigid tubes, is extended to address blood flow in deformable vessels where the elasticity of the vessel wall is significant. This theory is particularly useful for capturing wave propagation and phase lag effects in the cardiovascular system under pulsatile flow conditions, providing an analytical framework for understanding the interaction between blood flow and vessel wall motion (Alberto Figueroa et al. 2009; Filonova et al. 2020).

#### A.1 Governing Equations and Assump1ons

The following assumptions simplify the analysis:

1. **Axisymmetric Flow**: The flow and wall deformation are considered axisymmetric, meaning there is no dependency on the circumferential direction.
2. **Thin-Walled Elastic Vessel**: The vessel wall is assumed to be thin compared to its radius and behaves as an elastic material, allowing it to deform radially in response to pressure changes.
3. **Small Deformations**: The radial displacement of the wall is small, ensuring linearity in the elastic response.
4. **Oscillatory Flow**: The model assumes a periodic pressure gradient that drives pulsatile blood flow, resulting in oscillatory velocity and pressure fields.

Under these assumptions, we consider the Navier-Stokes equations for incompressible flow in cylindrical coordinates and couple them with equations describing the elastic deformation of the vessel wall.

#### A.2 Key Parameters

1. **Womersley Number α**:

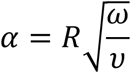

where, R is the vessel radius, ω=2πf is the angular frequency of the oscillatory pressure wave (with f being the frequency), ν is the kinematic viscosity of blood. The Womersley number characterizes the ratio of inertial to viscous forces. A high α\alphaα indicates that inertial effects dominate, leading to a parabolic velocity profile that flattens in the center of the vessel.
2. **Radial Displacement ξ(r**,**t):** The vessel wall undergoes radial displacement due to the pulsatile pressure. This displacement is governed by the wall’s material properties, specifically the Young’s modulus E, wall thickness h, and vessel radius R.

#### A.3 Formula1on of the Problem

1. **Navier-Stokes Equations** for Pulsatile Flow: The axial velocity w(r,t) of blood in a cylindrical vessel is described by:

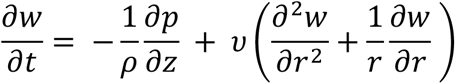

where, p(z,t) is the pressure, ρ is the density of blood, ν is the kinematic viscosity.
2. **Pressure Gradient**: The pressure p(z,t) is assumed to have an oscillatory component, typically given by:

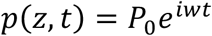

where *P*_0_ is the amplitude of the pressure wave, and ω is the angular frequency.
3. **Wall Deformation (Radial Displacement)**: The radial displacement ξ(r,t) of the wall is proportional to the oscillatory pressure applied on the inner wall. The relationship between radial displacement and wall pressure can be derived from the linear elasticity theory as:

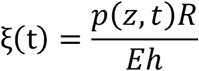

where, E is the Young’s modulus of the vessel wall, h is the wall thickness, R is the vessel radius.
4. **Velocity Profile**: The solution to the Navier-Stokes equation under these assumptions yields a velocity profile in the axial direction, which varies radially and temporally. The complex form of the velocity profile w(r,t) is:

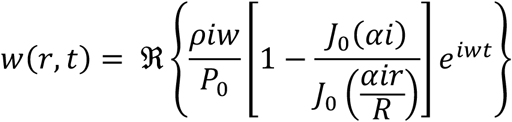

where *J*_0_ is the zeroth-order Bessel function of the first kind, capturing the radial dependence of the velocity profile.
5. **Circumferential and Axial Stresses**: In the elastic wall, the circumferential *σ*_*θ*_ and axial stresses *σ*_*z*_ are influenced by the pressure-induced deformation:

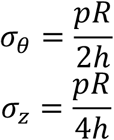

where *σ*_*θ*_ reflects the hoop stress, and *σ*_*z*_ represents the longitudinal stress in the vessel wall.

#### A.4 Analytical Solu1on for Pulsa1le Flow

The analytical solution provides insight into the phase difference between the pressure wave and the resulting velocity profile due to inertial effects and vessel compliance. For high Womersley numbers, the centerline velocity lags behind the pressure gradient due to the dominance of inertial forces, while for low Womersley numbers, the velocity profile follows a parabolic shape with minimal phase lag.

The total flow rate Q(t) through the vessel can be obtained by integrating the velocity profile across the cross-section:

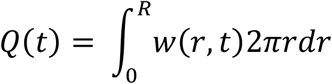

This flow rate is phase-shifted relative to the driving pressure wave, a key feature in understanding blood flow dynamics in compliant vessels.

#### A.5 Applications and Importance

Womersley’s deformable wall model has significant applications in cardiovascular modeling, particularly for assessing arterial compliance and wave propagation in large arteries. It is frequently used as a benchmark for verifying computational fluid-structure interaction (FSI) models, where the accuracy of blood flow simulation depends on accurately capturing vessel wall deformation under pulsatile loading.

The theoretical framework provided by Womersley’s model helps to characterize key hemodynamic parameters like wave speed, flow pulsatility, and vascular impedance, which are crucial for understanding conditions like arterial stiffness and hypertension.

### B. Constrained Mixture Model in a Single Vessel

A single segment of the arterial tree is considered as a thin-walled cylindrical tube composed of three main load-bearing constituents: elastin (*el*), collagen (*col*), and smooth muscle cells (SMCs; *smc*). First, we only consider the passive response of constituents. Each constituent is assumed to separately contribute to the strain energy density:

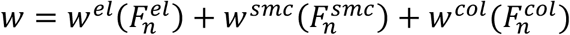

where 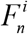 is the deformation gradient of each constituent *i* ∈ {*el, smc, col*} corresponding to its map from a stress-free configuration to the overall slow-time configuration. We define this deformation gradient as 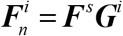, where ***G***^*i*^ is a pre-stretch for each constituent mapping each constituent from its distinct stress-free configuration to the intermediate configuration (Baek et al. 2005). In particular, the pre-stretch mapping for elastin can be expressed as

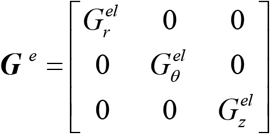

where 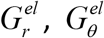, and 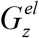 are pre-stretches associated with circumferential and axial directions, and 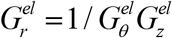. Similarly, for collagen fibers and SMCs, ***M***^***i***^ *i* ∈ {*smc*,*col*}, is defined as the unit vector in the direction of the collagen fiber (*k*) or SMCs. The pre-stretch mappings for collagen and smooth muscle cells are given as

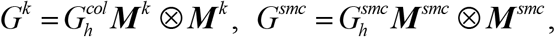

where the pre-stretches 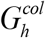 and 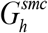 are also called homeostatic stretches, the stretches of the constituents when they are produced. We should note that in the previous applications of growth and remodeling, the pre-stretches were assumed to be constant for a single vessel. In our generalization of the framework to a vascular tree, we account for the variation of pre-stretches across the generations of vessels. Nevertheless, the pre-stretch implies that the homeostatic state in an individual vessel is associated with a constant homeostatic stress for the constituents of the vessel wall.

The orientation of collagen fibers and smooth muscles with respect to the axial direction in their reference configuration, defined by angle γ^*k*^, can be written as

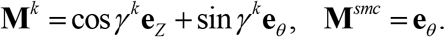

For modeling the extension and inflation of a thin wall model, ***F***^*s*^ is considered as ***F***^*s*^ = *diag*[*λ*_*r*_, *λ*_*θ*_, *λ*_*z*_]. The stretch in each constituent λ^*i*^ is expressed in terms of the prestretches using **F**^*i*^ *=* **F**^*s*^ **G**^*i*^:

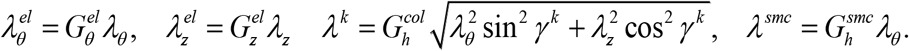

The incompressibility of the wall material is imposed by assuming an isochoric motion (i.e., **F**^*s*^ *=* 1, and thus *λ*_*r*_, *= (λ*_*θ*_, *λ*_*z*_)^−1^. Using the membrane theory (Humphrey 2002), the membrane Cauchy stress (force per deformed length) can be written as

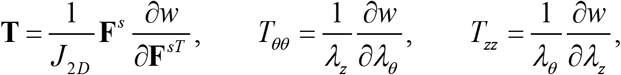

where *J*_*2D*_ = *λ*_*θ*_, *λ*_*z*_ The total strain energy per unit area can be written as

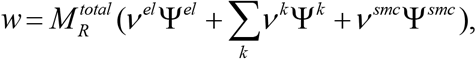

where *v*^*el*^, *v*^*k*^, and *v*^*smc*^ are mass fractions of elastin, collagen fiber families, and SMCs, respectively. In this work, four families of collagen fibers in circumferential, axial, and two diagonal directions were considered with mass fractions *v*^*K*^ *=* (0.1,0.1,0.4,0.4)*V*^*col*^ where *V*^*col*^ is the total collagen mass fraction(Zeinali-Davarani et al. 2011). The total mass per unit area 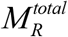 is the mass of load bearing constituents and can be computed via

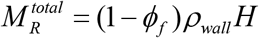

where *ρ*_*wall*_ is the density of the vascular wall, *ϕf* is the volume fraction of interstitial fluid, and *H* is the wall thickness under homeostatic conditions.

A neo-Hookean model is employed for the passive elastin response and a Holzapfel exponential model is used for collagen fiber families and passive behavior of circumferentially oriented SMCs

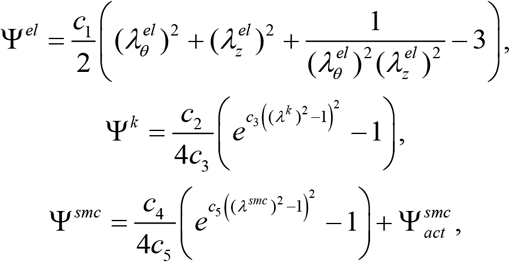

where *c*_1_ is the elastin material parameter, *c*_2_ and *c*_3_ are collagen material parameters, and *c*_4_ and *c*_5_ are passive SMC material parameters. To include the active tension of vascular SMCs, we use a potential function as given by (Baek et al. 2007)

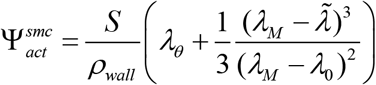

where *λ*_*M*_ and *λ*_0_ are stretches at which the active force generation is maximum and zero, respectively, and *S* is the basal active tone. In addition, 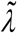 is an active stretch of the SMCs in the circumferential direction, which can evolve by SMC remodeling over slow Timescales (hours to days). In the current study, we assume 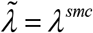. Finally, for pressurized thin-walled cylinder with mid-vessel pressure law 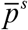, the force equilibrium in the circumferential direction gives the Laplace

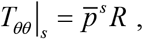

where *R* is the vessel radius. In our homeostatic optimization framework, we assume the intermediate configuration of the blood vessel in our continuum mechanics formulation is its homeostatic configuration. Therefore, by setting *F*^*s*^ = (i.e., *λ*_*θ*_ *= λ*_*z*_ = 1)in equations above, the homeostatic membrane stress **T**^*i*^ and total stress, 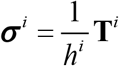 can be determined as a function the passive material properties (*c*_1_ − *c*_5_), constituent prestretches 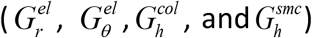 and *!* their mass fractions *v*^*i*^, and active SMC parameters, *λ*_*M*_, *λ*_0_ and *S*.

### C. Slow-time hemodynamics

Here we present an algorithm that computes the slow-Time pressure and flow at each vessel for a given bifurcating tree. We introduce the index [*k*,*m*] to label an individual vessel within the tree generation *k* for *m =* 1,..2^*kࢤ1*^. A large sparse matrix can be constructed to account for the pressure continuity and flow conservation at each bifurcation as well as Poiseuille flow resistance at each vessel. The block-matrix, related to the vessel [*k*,*m*], comprises of flow splits and Poiseuille equation for daughter vessels in generation *k* +1, as shown below

**Given resistance** *res*[*k*,*m*]:

**Construct** [*k*,*m*] **– block of equations:**

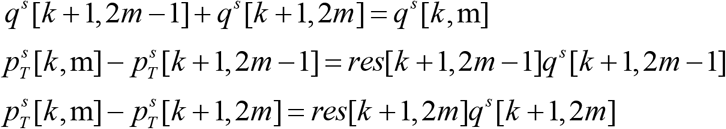

**for unknowns:**

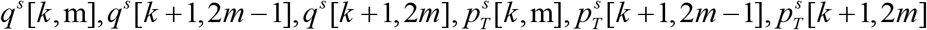

For boundary conditions in the slow-time hemodynamic system in a tree, we consider two options: (1) given input steady flow at the root vessel and terminal steady pressure at outlets; and (2) given input steady pressure at root vessel and steady flow at outlets. In the manuscript, for the numerical example of the symmetric tree we consider boundary conditions (1) while for the asymmetric tree example (2).

### D. Fast-time Hemodynamics

Here we present an algorithm that for a given geometry of a bifurcating tree and discrete frequency *ω*_*n*_ computes the impedance and then fast-time hemodynamics at each vessel. First, we use the following relations between characteristic *Z*^*c*^ terminal *Z*^*T*^, and input *Z*^*inp*^ impedance of each vessel, and the reflection coefficient

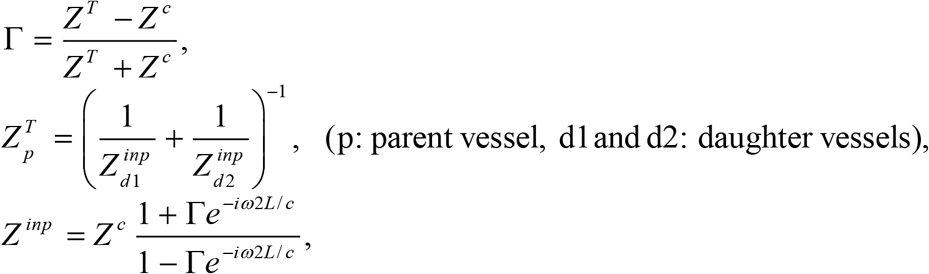

where *L* and *c* are vessel length and wave velocity, respectively. First, we compute the impedance at each vessel from bottom-to-top of the tree. We consider that the terminal impedance or, equivalently, the reflection coefficients Γ at outlets are given

**Given:** 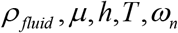

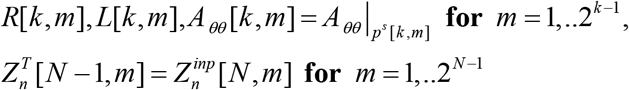

**Find:** 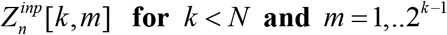

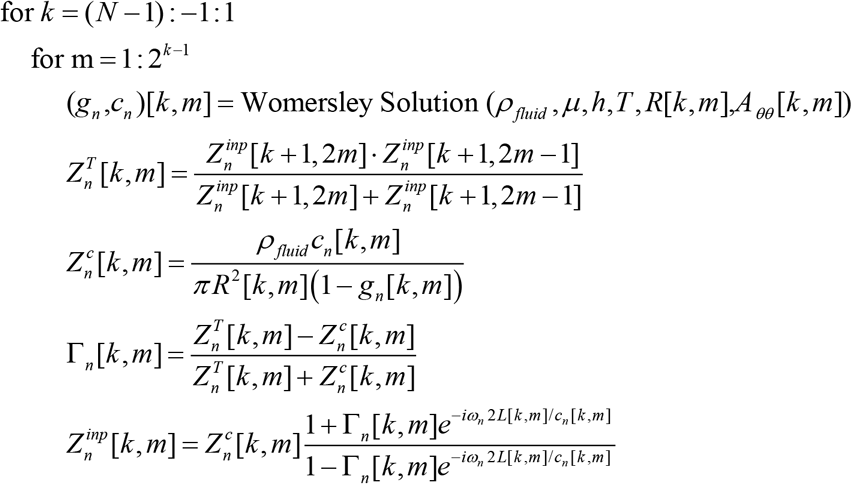

end

end

Second, we reconstruct the pulsatile hemodynamics in the system by computing the pressure and flow solution at each vessel from top-to-bottom of the tree. The components of the fast-time terminal pressure 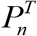 and input flow 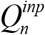 in frequency domain are found using from Womersley’s solution. For illustration of the algorithm, we consider the boundary condition at the root vessel as a given input pressure.

**Given:** 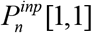

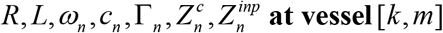

**Find:** 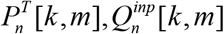

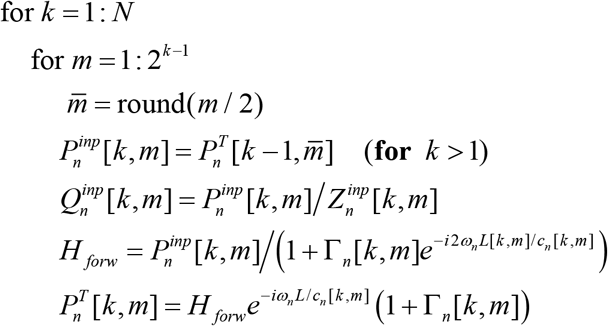

end

end

